# BioSharing: Harnessing Metadata Standards for the Data Commons

**DOI:** 10.1101/144147

**Authors:** Susanna-Assunta Sansone, Alejandra Gonzalez-Beltran, Philippe Rocca-Serra, Peter McQuilton, Massimiliano Izzo, Allyson Lister, Milo Thurston

## Abstract

The use of community-driven metadata standards, such as minimal information guidelines, terminologies, formats/models, is essential to ensure that data and other digital research outputs are Findable, Accessible, Interoperable, and Reusable, according to the FAIR principles. As with other types of digital assets, metadata standards also need be FAIR. Their discoverability and accessibility is ensured by BioSharing, the most comprehensive resource of metadata standards, interlinked to data repositories and policies, available in the life, environmental and biomedical sciences. With its growing content, endorsements, and collaborative network, BioSharing is part of a larger ecosystem of interoperable resources. Here we describe some of the activities under the USA National Institutes of Health (NIH)’s Big Data to Knowledge (BD2K) Initiative, illustrating how we track the evolution and use of metadata standards and work to connect them to indexes and annotation tools.

## BIOSHARING: AN INFORMATIVE AND EDUCATIONAL RESOURCE

In the life, environmental and biomedical sciences there are almost a thousand community-developed content standards - encompassing reporting guidelines (or checklists), models/formats and terminologies (e.g. taxonomies, ontologies) - many of which have been created and/or implemented by several thousand data repositories or databases. Content or metadata standards ensure that the relevant elements of a dataset (e.g., fundamental biological entities or experimental components, as well as complex concepts such as tissues and diseases, along with the analytical process and the mathematical models) are reported consistently and meaningfully, opening the datasets to transparent interpretation, verification, exchange, integrative analysis and comparison, supporting the FAIR principles[1]. Along with other community-developed guidelines, such as those on data citation[2] and identification[3], metadata standards are designed to assist the virtuous data cycle, from collection to annotation, through preservation and publication to subsequent sharing and reuse. A recent Wellcome Trust-commissioned review[4] provides an overview on the variety of interoperability standards, focussing on metadata standards, their role in research data management and the related challenges and opportunities.

For the consumers of these metadata standards, it is often difficult to know which are the most relevant for a specific domain or need; while for producers it is important that their resources are findable by prospective users[5]. These are the key use cases addressed by BioSharing[6,7], a curated, informative and educational resource with over 1,700 records describing and interlinking metadata standards, data repositories, and policies in the life, environmental and biomedical sciences. Specifically, BioSharing works with and for the community to map the landscape of metadata standards, defining the indicators necessary to monitor their evolution, implementation and use in data repositories, and their adoption in data policies by funders, journals and other organizations.

Launched in 2011 as an evolution of the MIBBI portal[8], BioSharing is driven by an international advisory board and reaches out to researchers, developers, curators, funders, journal editors, librarians and data managers worldwide[9], through a joint Force11[10] and the Research Data Alliance (RDA)[11] working group. BioSharing has already been adopted by a variety of communities and stakeholders, including publishers (e.g. Springer Nature, PloS, EMBO press and Wellcome Trust Open Research), standardization groups, research data management support initiatives and libraries. BioSharing has also been endorsed by the European ELIXIR programme[12], and its relevance highlighted in reports from two workshops by the USA National Institutes of Health (NIH)’s Big Data to Knowledge (BD2K) Initiative[13,14,15].

## EXAMPLES OF USE AND ACTIVITIES

We describe three examples of how BioSharing contributes to the NIH BD2K community and some projects, as part of ongoing research and development activities. These exemplars are quite diverse and allow us to show some of the existing BioSharing features and future directions, serving both researchers and developers and striving to embed metadata standards into the data cycle in an ‘invisible’ manner.

### Tracking the evolution of metadata standards

BioSharing content can be searched using simple or advanced searches, refined via our filtering options, or grouped via the ‘Collection’ feature, according to field of interest or focus. For example, journals and publishers are collating the metadata standards and data repositories they recommend in their data policies. Similarly, communities, projects and organizations are creating Collections by selecting and filtering standards (and data repositories) relevant to their work, and/or those they are actively developing themselves. An example of the latter case, is provided by the NIH Library of Network-Based Cellular Signatures (LINCS) Program[16], which is creating a network-based understanding of biology by cataloging changes in gene expression and other cellular processes that occur when cells are exposed to a variety of perturbing agents.

As part of this multi-site program, the LINCS Data Working Group (DWG) works to develop metadata standards to describe LINCS reagents, assays and experiments[17] to ensure that key elements of experimental metadata are reported in a common manner, facilitating the metaanalysis between all LINCS Centers and the release of FAIR data to the community via the LINCS Data Portal. Due to the dynamic nature of the experiments and the phased development of the standards specifications, the LINCS DWG uses BioSharing to display and track the evolution of their metadata standards.

BioSharing has enabled the LINCS DWG to create, edit and maintain their own records for their standards, and group them under a LINCS Collection[18]. Each record can have one or more maintainer, who has a user profile that can be linked to their resources, publications and ORCID identifier[19]; this provides not only visibility for the individuals but also a much needed contact point for prospective users. Each metadata standard (and data repository) record in BioSharing is manually curated to ensure its description and status is up-to-date, and the validity of the information therein is checked with the maintainers and/or the community behind each effort. Figure 1 illustrates an example of how the BioSharing indicators (of readiness for implementation or use) are used to show the evolution of one of the LINCS standards.

**Figure 1.**
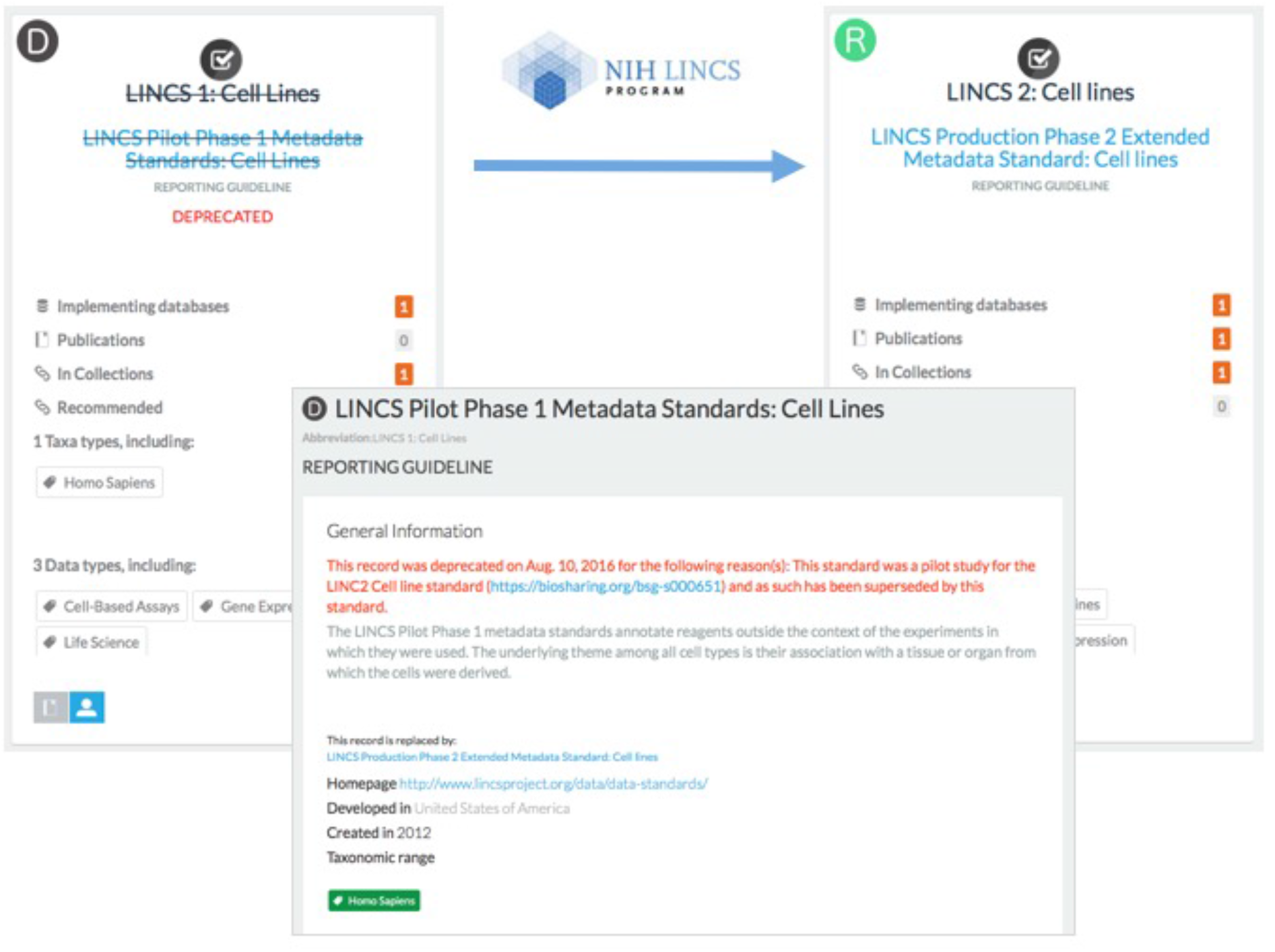
An example of two metadata standards, specifically reporting guidelines: ‘LINCS 1: Cell Lines’ (tagged with ‘D’, indicator for deprecation) and has been superseded by ‘LINCS 2: Cell Lines’ (tagged with ‘R’, indicator for ready for use). When available, the reason for the deprecation is detailed in the deprecated record.

### Linking data repositories to relevant standards

Another example of a Collection is the one created with and for the the NIH BD2K bioCADDIE project. The bioCADDIE Collection[20] groups and displays the existing metadata standards that have been used to develop the DatA Tag Suite (DATS), the metadata model[21] underpinning DataMed[22]. This data index and search engine prototype, is based on metadata extracted from various data sets in a range of data repositories, and does for data what PubMed[23] has done for the literature.

One of the search use cases elicited from researchers during the DataMed development phase, is to allow the searching and filtering of datasets (from data repositories) that are compliant with a given community metadata standard. For this reason the DATS model is designed around the *Dataset* metadata element that is linked to other digital objects, such as *Publication, Software, DataRepository* and *DataStandard*, which are the focus of other indexes, such as PubMed, BD2K Aztec[24] and BioSharing, respectively. The latter link is especially important, because knowing if a data repository uses open community standards to harmonize the reporting of its different datasets will provide researchers with some confidence that these datasets are (in principle) more comprehensible and reusable.

Figure 2 shows an example of how BioSharing builds this interlinkage between metadata standards and the data repositories that implement them, as well as showing their indicators of readiness. BioSharing is therefore well placed to provide DataMed with the knowledge of the relation(s) between metadata standards and the data repositories that implement them. To realize this query, work is in progress to deliver a BioSharing ‘look up service’ functionality that will allow systems like DataMed to access and use the information in the context of their searches.

**Figure 2.**
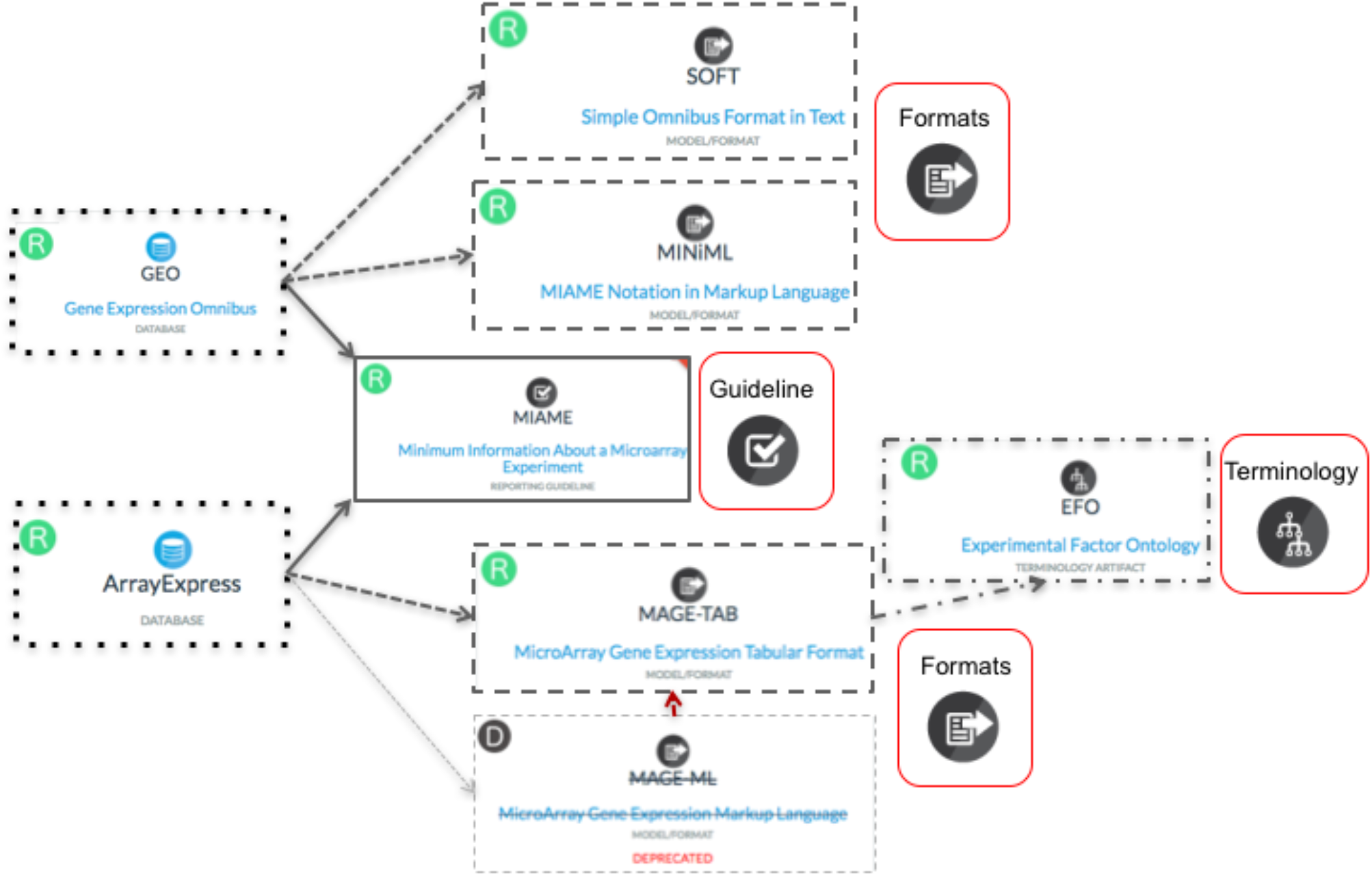
An example of interlinking between metadata standards (reporting guidelines, models/formats and terminologies) and data repositories that implement them, along with their indicators of readiness. This shows how two data repositories of transcriptomics data implement the same reporting guideline, but use different formats for the upload/download of the datasets with only one of the repositories implementing a standard terminology (to describe experimental factors).

### Driving annotation and validation against metadata standards

Despite the growing set of reporting guidelines, models/formats and terminologies for describing the experiments, the barriers to authoring the experimental metadata necessary for sharing and interpreting datasets are still tremendously high. The reasons are twofold. First, bound by a particular discipline or domain, metadata standards are fragmented, with gaps and duplications, thereby limiting their combined usage. For example, producers of datasets in which source material has been subjected to several kinds of analysis (e.g., genomic sequencing and clinical measurement) find it particularly challenging to describe the datasets as coherent units of research due to the diversity of metadata standards with which the parts must be formally represented. Second, understanding how to comply with these metadata standards takes time and effort, and researchers often see them as burdensome and/or over-prescriptive, as something that may benefit other scientists, but not themselves. In addition, these guidelines are usually narrative in form and prone to be ambiguous, further making their compliance difficult.

The need for tools and services that facilitate the ‘invisible use’ of metadata standards is widely recognized; this is something of which we have first-hand experience via the ISA framework[25, 26]. Further research is ongoing in BioSharing to explore how to automatically use metadata requirements, from two or more domain specific standards, for composition in annotation templates and for validation purposes. To this end, BioSharing contributes to the NIH BD2K CEDAR project[27, 28], which works to develop tools and practices to make the authoring of complete datasets smarter and faster.

Figure 3 shows how BioSharing works to define the method and process to create modularized metadata elements, tracking provenance (e.g., information about the standard(s) the metadata elements are derived from, as well as the process of derivation), conditions and dependencies (of each standard(s)-derived metadata element) and validation rules (to ensure a template meets the requirement of one or more checklists). Ultimately, the machine-readable versions of the standards-derived metadata elements will be served to inform the creation of descriptive templates in the CEDAR, and/or validation of datasets in others tools like ISA.

**Figure 3.**
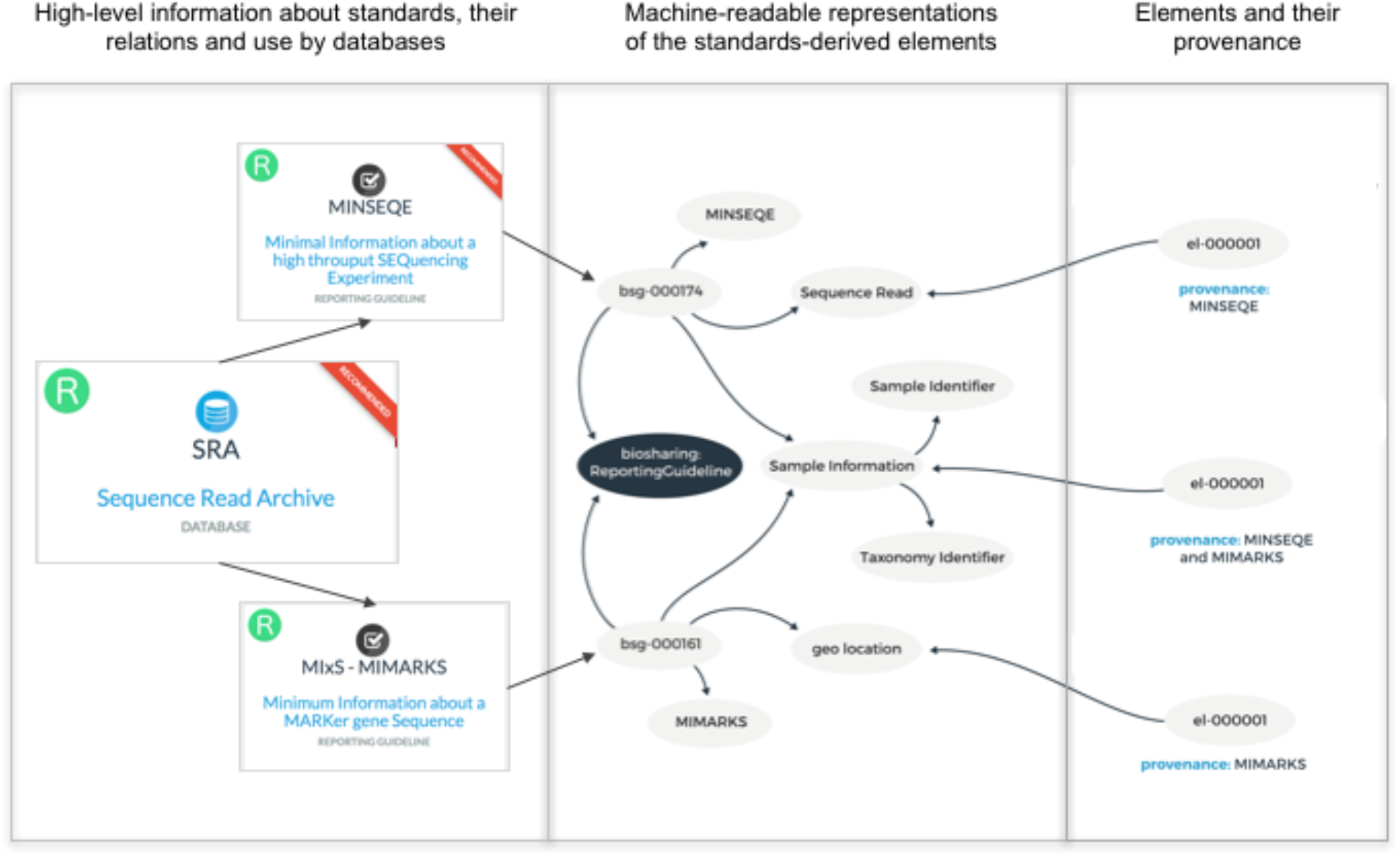
An example of machine-readable metadata elements, and their provenance information, produced from community-defined metadata standards (required by some data repositories). Specifically, two reporting requirements implemented by the Sequence Read Archive (SRA) to describe generic (MINSEQE) and environmental survey-based (MiXS-MIMARKS) sequencing experiments; the red ‘Recommended’ banner shows that SRA and MINSEQE are recommended by one or more publishers’ data policies.

## BIOSHARING: AN ELEMENT OF THE COMMONS

To implement the FAIR principles it is necessary to: (i) have a comprehensive description of standards; and (ii) help researchers, developers, curators, funders, journals and librarians to best navigate and select the various standards, or to find the repositories that implement them and draft a data management plan; or simply to find enough information to make an informed decision on which standards or related repository should be recommended in a data policy, funded or implemented.

Via its informative and educational functionalities and indicators, BioSharing provides: (i) developers of standards and data repositories with a mean to increase the discoverability of their resources outside their own direct community, and (ii) prospective consumers with ways to visualize and understand the status of these resources, enabling them to make an informed decision as to which standard (database or policy) to (re)use or endorse, thus maximizing the potential of adoption and reducing the potential for unnecessary reinvention.

A recently conducted survey[29], supported by ELIXIR and NIH BD2K, provides an insight of users’ needs from BioSharing, showing the road ahead and driving our future activities. As highlighted by the Wellcome Trust-commissioned review[4], it is essential to recognize interoperability standards as digital objects in their own right, with their associated research, development and educational activities.

## ACKNOWLEDGEMENTS

We thank all contributors to the BioSharing project, our user community, the Advisory Board Members and the members of the RDA and Force11 Working Groups. We also thank our ELIXIR and BD2K colleagues, in particular those part of the ELIXIR Interoperability Platform, the bioCADDIE consortium, the CEDAR project and the LINCS programme.

## REFERENCE LIST

1. Wilkinson MD, Dumontier M, Aalbersberg IJ, et al. The FAIR Guiding Principles for scientific data management and stewardship. Sci Data. 2016;15(3):160018

2. Fenner M, Crosas M, Grethe J, et al. A Data Citation Roadmap for Scholarly Data Repositories. Preprint bioRxiv 097196; doi: https://doi.org/10.1101/097196, accessed May 2017.

3. McMurry J, Juty N, Blomberg N, et al. Identifiers for the 21st century: How to design, provision, and reuse persistent identifiers to maximize utility and impact of life science data. Preprint bioRxiv 117812; doi: https://doi.org/10.1101/117812, accessed May 2017.

4. Sansone SA and Rocca-Serra P. Interoperability Standards - Digital Objects in Their Own Right. Wellcome Trust. 2016. https://doi.org/10.6084/m9.figshare.4055496.v1, accessed May 2017.

5. Tenenbaum JD, Sansone SA, Haendel M. A sea of standards for omics data: sink or swim? J Am Med Inform Assoc. 2014;21(2):200–3.

6. McQuilton P, Gonzalez-Beltran A, Rocca-Serra P, et al. BioSharing: curated and crowd-sourced metadata standards, databases and data policies in the life sciences. Database (Oxford). 2016;2016. pii: baw075.

7. BioSharing portal: https://biosharing.org, accessed May 2017.

8. Taylor CF, Field D, Sansone SA, et al. Promoting coherent minimum reporting guidelines for biological and biomedical investigations: the MIBBI project. Nat Biotechnol. 2008;26(8):889–96.

9. BioSharing adopters and collaborators: https://biosharing.org/communities, accessed May 2017.

10. BioSharing working group page under Force11: https://www.force11.org/group/biosharingwg, accessed May 2017.

11. BioSharing working group page and recommendation under RDA: https://www.rd-alliance.org/group/biosharing-registry-connecting-data-policies-standards-databases-life-sciences.html, accessed May 2017.

12. ELIXIR website: https://www.elixir-europe.org, accessed May 2017.

13. Executive Summary - Workshop on Frameworks for Community-Based Standards Effort Workshop” NIH BD2K, Sep 2013: https://datascience.nih.gov/sites/default/files/bd2k/docs/frameworksforcommbasedstandardseffortsreport.pdf, accessed May 2017.

14. Sansone, SA, Derr KL, Kennedy KD and Huerta M. NIH BD2K workshop report: Frameworks for Community-based Standards Efforts. Figshare. https://doi.org/10.6084/m9.figshare.3795816.v2, accessed May 2017.

15. Executive Summary - Workshop on Community-based Data and Metadata Standards Development: Best practices to support health y development and maximize impact” NIH BD2K, Feb 2015: https://datascience.nih.gov/sites/default/files/bd2k/docs/ExecSummCBDMSworkshopFEB2015.pdf, accessed May 2017.

16. LINCS programme website: http://www.lincsproject.org, accessed May 2017.

17. LINCS data standards website: http://www.lincsproject.org/LINCS/data/standards, accessed May 2017.

18. LINCS Collection in BioSharing: https://biosharing.org/collection/LINCSProject, accessed May 2017.

19. ORCID website: https://orcid.org, accessed May 2017.

20. BioCADDIE Collection in BioSharing: https://biosharing.org/collection/bioCADDIE, accessed May 2017.

21. Sansone, SA, Gonzalez-Beltran A, Rocca-Serra P, et al. DATS: the data tag suite to enable discoverability of datasets. Sci. Data 2017;4:170059.

22. Ohno-Machado L, Sansone SA, Alter G, et al. Finding useful data across multiple biomedical data repositories using DataMed. Nat Genet. 2017;49(6):816–819.

23. PubMed website: https://www.ncbi.nlm.nih.gov/pubmed, accessed May 2017.

24. Aztec website: https://aztec.bio, accessed May 2017.

25. ISA framework website: http://isa-tools.org, accessed May 2017.

26. Sansone SA, Rocca-Serra P, Field D, et al. Toward interoperable bioscience data. Nat Genet. 2012;44(2):121–6.

27. Musen MA, Bean CA, Cheung KH, et al. The center for expanded data annotation and retrieval. J Am Med Inform Assoc. 2015;22(6):1148–52.

28. CEDAR website: https://metadatacenter.org, accessed May 2017.

29. BioSharing survey summary report: https://doi.org/10.6084/m9.figshare.3795810.v2, accessed May 2017.

